# Cholesteryl Ester as a Prognostic Biomarker In IDH-wildtype Glioblastoma

**DOI:** 10.64898/2026.05.05.722825

**Authors:** Nana Wang, Jiejun Wang, Jianlin Liu, Jinqi Zou, Bin Yang, Pu Wang, Nan Ji, Shuhua Yue

**Affiliations:** Key Laboratory of Biomechanics and Mechanobiology (Beihang University), Ministry of Education; Key Laboratory of Innovation and Transformation of Advanced Medical Devices, Ministry of Industry and Information Technology; National Medical Innovation Platform for Industry-Education Integration in Advanced Medical Devices (Interdiscipline of Medicine and Engineering); School of Biological Science and Medical Engineering, Beihang University, Beijing, 100191, China; Department of Neurosurgery, Beijing Tiantan Hospital, Capital Medical University, Beijing, 100070, China; Department of Cerebrovascular Diseases, Beijing Anzhen Hospital, Capital Medical University, Beijing, 101118, China; Beijing Wellmed Medical Diagnostics & Laboratory Co., Ltd, Beijing, 102600, China

**Author notes:** Corresponding authorship. (S.Y.); (N.J.). Equal contribution (Nana Wang, Jiejun Wang).

**Keywords:** Biomarker, cholesteryl ester, IDH-wildtype GBM, stimulated Raman scattering, prognostic model

## Abstract

Current treatment of IDH-wildtype glioblastoma (GBM) relies on the first-line chemotherapy-temozolomide. Although MGMT methylation is routinely conducted to predict chemosensitivity, its efficacy is often compromised. Thus, there is an urgent need to discover more accurate prognostic biomarkers. Cholesteryl ester (CE) has been recently recognized as a key feature of GBM, however, its role in GBM prognosis remains poorly understood. We first employed label-free stimulated Raman scattering (SRS) imaging to quantitatively analyze CE level in intact tumor tissues obtained from IDH-wildtype GBM patients. Our result revealed significantly prolonged 2-year overall survival (OS) in patients with CE level ≥ 40% compared to those with CE level < 40%. CE outperformed MGMT methylation for 2-year OS prognosis (AUC: 0.836 vs. 0.763). Importantly, CE also achieved superior prognostic performance over MGMT methylation on an independent cohort, with higher sensitivity (0.856 vs. 0.667), specificity (0.833 vs. 0.583), NPV (1.00 vs. 0.667), PPV (0.833 vs. 0.583). Given synergistic effects between CE and MGMT methylation, we developed a prognostic model combining these two biomarkers. Specially, machine learning (XGBoost) model exhibited optimal performance in the training cohort (AUC: 0.920), and maintained its superior performance on the independent cohort (sensitivity: 0.946, specificity: 0.873, NPV: 1.00; PPV: 0.917). Mechanistically, integrative analysis of TCGA database linked poor prognosis to the coordinated upregulation of genes involved in cholesterol efflux, hydrolysis, transport, and inhibition of de novo synthesis, unraveling a possible underlying mechanism between poor prognosis and cholesterol metabolism. This work identified CE as a prognostic biomarker for IDH-wildtype GBM.

## INTRODUCTION

The 2016 World Health Organization (WHO) classification of gliomas underscores the prognostic significance of isocitrate dehydrogenase (IDH).^1^ Specifically, approximately 90% of glioblastoma (GBM) are IDH-wildtype and exhibit poorer prognosis compared to IDH-mutant ones.^2^ The 2021 WHO classification further refined the categorization based on molecular features, redefining IDH-mutant gliomas as astrocytoma, IDH-mutant, WHO 4, while IDH-wildtype gliomas retain the designation of GBM.^3,4^ Notably, IDH-wildtype GBM is the most prevalent and aggressive glioma subtype in adults. Current standard of care for GBM includes maximal safe resection, adjuvant radiation, and chemotherapy with the first-line alkylating agent temozolomide (TMZ).^5–7^ Chemosensitivity to TMZ primarily associates with epigenetic silencing by methylation of the O (6)-Methylguanine-DNA methyltransferase (MGMT) methylation.^8–11^ While MGMT methylation remains a key predictor of chemosensitivity, its efficacy is often compromised by several challenges. These include non-quantitative methods, cumbersome procedure, prolonged detection times, large variation of cutoff value, or cost-intensive analysis.^12–15^ Specially, the diagnostic performance of current assays (as measured by the area under the ROC curve, AUC) typically falls below 0.8 in clinical validation studies.^16–18^ These challenges underscore the urgent need to identify novel biomarkers that complement or surpass MGMT methylation, enabling more precise and adaptive approaches to IDH-wildtype GBM prognosis.

Metabolic reprogramming has been recognized as a key feature of GBM.^19–21^ For accommodating vigorous lipid turnover, excessive lipid is esterified into neutral lipid, primarily triacylglycerol (TAG) and cholesteryl ester (CE), and subsequently stored in lipid droplets (LDs).^22^ Previous studies have focused on the presence of CE in GBM and its potential as a therapeutic target.^23–25^ These studies encompassed patients harboring IDH-mutant GBM and therefore lag behind the 2021 WHO classification, which classify previously defined IDH-mutant GBM as astrocytoma, IDH-mutant, WHO 4.^3, 4^ Notably, emerging evidence suggests that CE level correlates with IDH status of gliomas, where IDH-wildtype GBMs display significantly lower CE level compared to IDH-mutant counterparts.^26^ Our recent work has unveiled remarkable CE level difference even within IDH-wildtype GBM.^27^ Thus, it is intriguing to explore whether CE could serve as a prognostic biomarker for IDH-wildtype GBM.

To assay for lipid composition, analytical tools, such as mass spectrometry and nuclear magnetic resonance spectroscopy, are commonly used. Because such techniques require big tissue homogenization, which conflicts with the clinical necessity to preserve precious specimens by minimizing tissue. Mass spectrometry imaging has been developed to map the spatial distribution of biomolecules in intact tissues, but could hardly observe submicron-scale LDs.^28^ Owing to the high molecular selectivity and submicron spatial resolution, stimulated Raman spectroscopy (SRS) has made pivotal contributions to the study of the role of LD in various types of cancer.^29–37^ It is conceivable that SRS imaging could provide new insights into the relationship between CE and prognosis in IDH-wildtype GBM.

Herein, we used label-free SRS imaging to quantitatively map the CE level in IDH-wildtype GBM tissues obtained from patients undergone resection surgery. Figure 1 showed the flowchart. First, utilizing SRS imaging, we achieved spatially resolved quantification of CE within LDs in IDH-wildtype GBM tissues. Our results revealed a significant survival benefit in patients with CE level ≥ 40% compared to the ones with CE level < 40%. Second, we found CE outperformed MGMT methylation in prognostic prediction. Leveraging synergistic effects between CE and MGMT methylation, we further used machine learning to develop a prognostic model based on these two biomarkers, and tested the model on an external cohort. Final, we elucidated the mechanistic link between cholesterol metabolism and prognosis using the cancer genome atlas (TCGA) database. These findings collectively develop a novel prognostic platform discovering CE as a superior biomarker and unravel the altered cholesterol metabolism in IDH-wildtype GBM with poor prognosis, which may open up new opportunities for the diagnosis and treatment of aggressive IDH-wildtype GBM.

**Figure 1.**
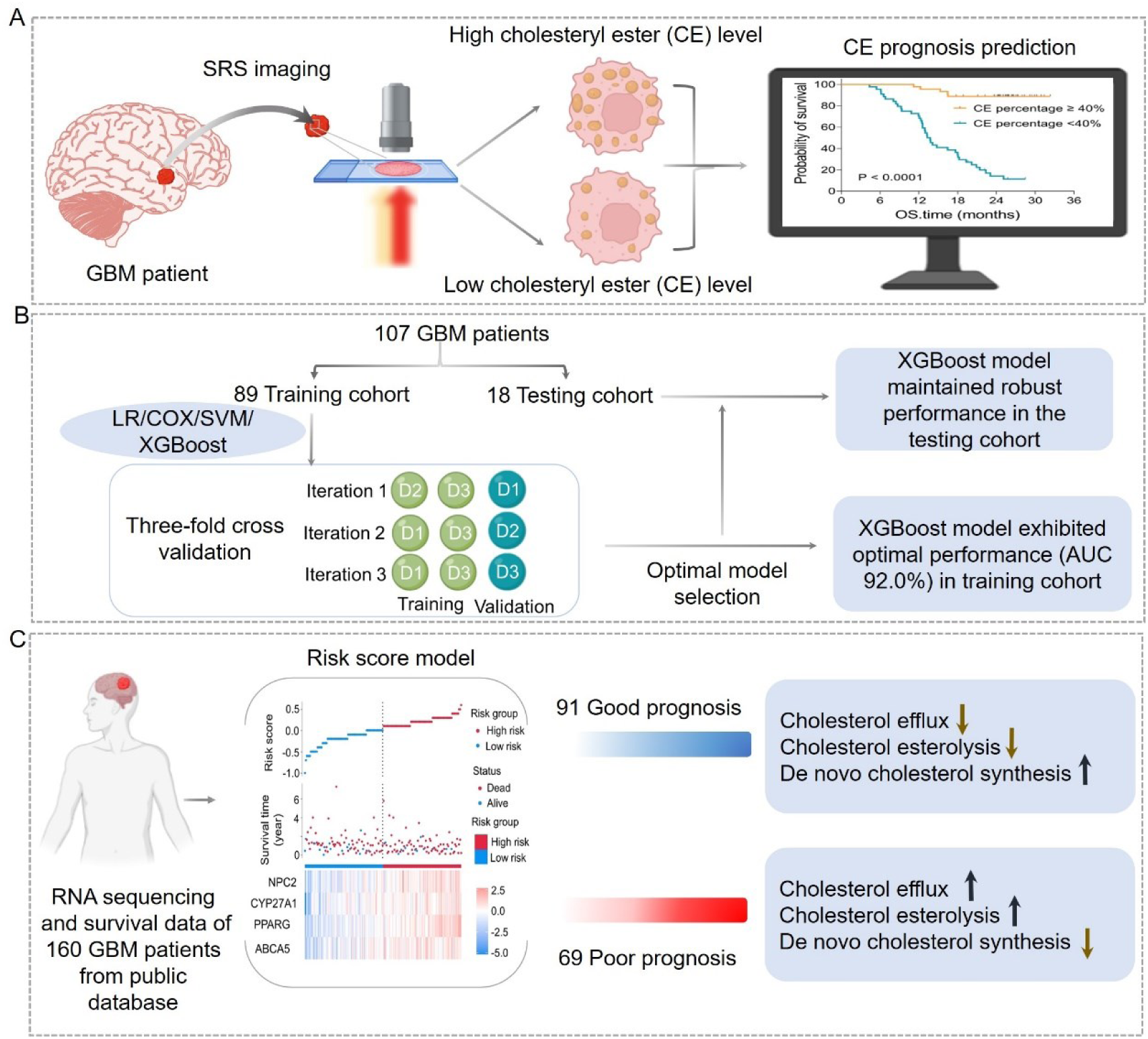
Overview of the workflow. (A) The label-free SRS imaging to quantitatively map the CE level in IDH-wildtype GBM tissues obtained from patients undergone resection surgery, revealing significant survival difference (*P* < 0.0001). (B) Machine learning framework addressing the limited cohort through three-fold cross-validation. Four distinct models (COX, LR, SVM, and XGBoost) underwent iterative training and validation cycle. XGBoost achieved optimal training accuracy. XGBoost further was teste in an independent cohort, and maintained robust prognostic performance. (C) The risk score model, trained on survival data from 160 patients in a public database, categorized 91 patients as favorable prognosis and 69 as poor prognosis. Subsequent analysis evaluated cholesterol metabolism dynamics, including efflux, hydrolysis, and de novo synthesis pathways.

## RESULTS

### Clinical Implications of CE in GBM Prognostication via SRS Imaging

To ensure homogeneity of the patient cohort (*e.g.* in terms of the IDH status) and clinical protocol standardization, we recruited a homogeneous group of 107 IDH-wildtype GBM patients with a Karnofsky Performance Status (KPS) > 70%, who received i) Stupp regime within 4-6 weeks after initial surgery, and ii) completed Stupp regime after 6 cycles or until progression of disease, assessed according to the Response Assessment in Neuro-Oncology (RANO) criteria. The cohort was further stratified into two groups based on a two-year OS. Patients with OS ≥ 2 years was high OS group and those with OS < 2 years was low OS group. All clinical information is provided in Table S1.

Using an SRS imaging-based quantitative platform, we analyzed CE level within LDs in IDH-wildtype GBM tissues. By tuning the laser beating frequency to be resonant with C-H stretching vibration at 2850 cm^-1^, strong SRS signals arose from the lipid-rich cell membranes and LDs, and in the meantime, weak signals were derived from the lipid poor nuclei. Such imaging contrast permits the clear visualization of cellular morphology and intracellular LDs in a label-free manner. As shown in Figure 2A, we found a substantial reduction in LD accumulation in low OS GBM compared to the high OS GBM. Further composition analysis of individual LDs by spontaneous Raman spectra identified high CE level, based on the distinctive Raman peaks for cholesterol ring around 702 cm^-1^ and ester bond around 1742 cm^-1^, in high OS GBM, but not in low OS GBM (Figure S1). The corresponding SRS spectra in the C-H vibration region (2800-3050 cm⁻¹) identified distinct cholesterol Raman band (2870 cm^-^¹) in high OS GBM, but not in low OS GBM (Figure 2B). To quantitatively map CE in LDs, we established a calibration curve for CE concentration using CE-TG mixture emulsions of different CE molar percentages (R² = 0.99, Figure S2), following previously reported method.^38,39^ The quantitative CE imaging was achieved on the basis of the calibration curve (Figure S3). As shown in Figure 2C, the representative CE imaging suggested that CE level was greater in high OS GBMs than in low OS ones. Quantitative analysis showed that high OS GBMs exhibited significantly higher CE level (48.76% ± 2.33%) compared to low OS ones (CE: 32.26% ± 2.08%; *P* < 0.0001, Figure 2D and Table S1). Collectively, these findings demonstrate that CE may be associated with the IDH-wildtype GBM prognosis.

**Figure 2.**
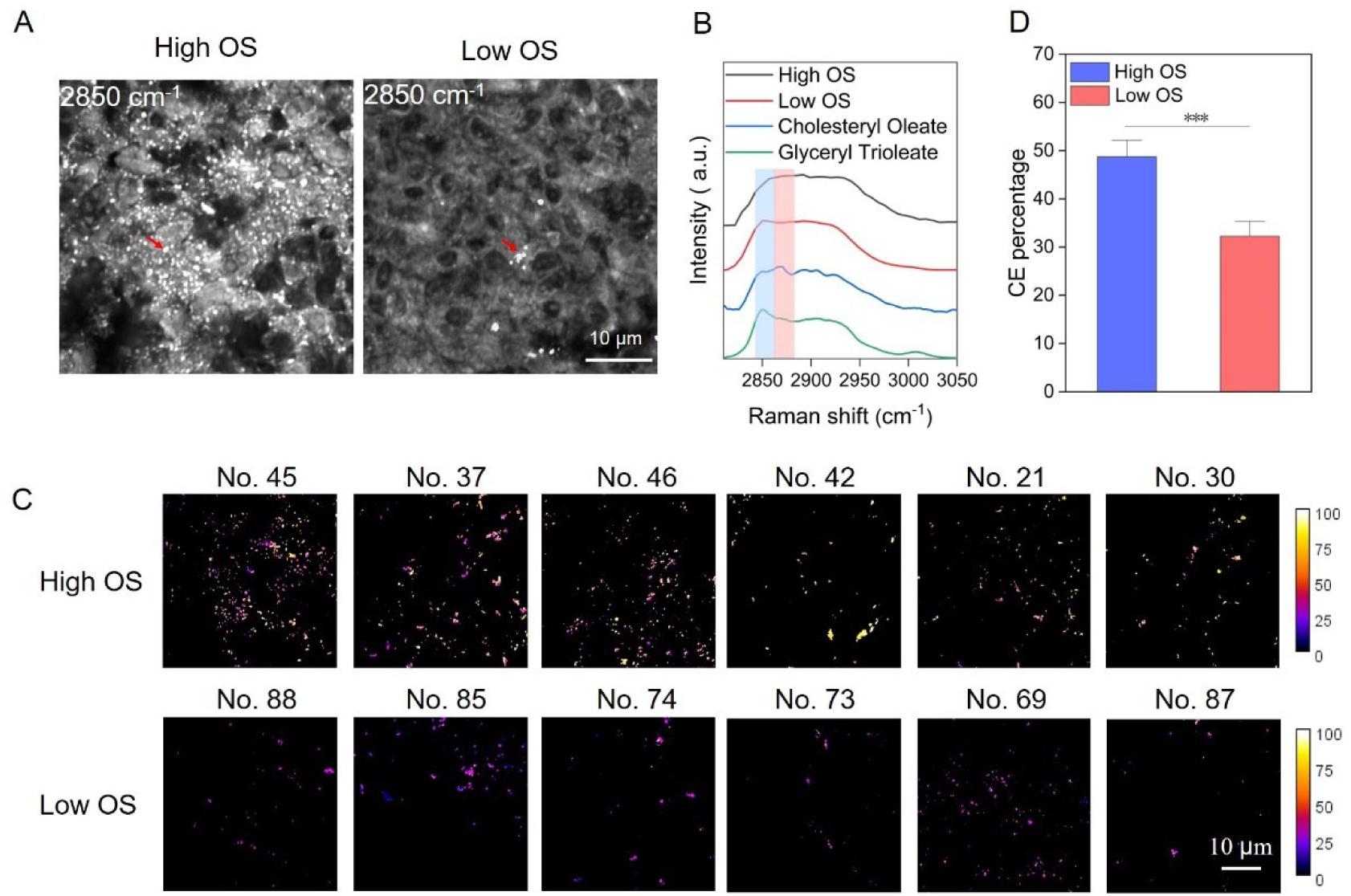
Label-free quantitative SRS imaging of CE in IDH-wildtype GBM tissues. (A) Representative SRS images of IDH-wildtype GBM tissues from high OS and low OS cohorts, acquired at 2850 cm⁻¹. LDs are indicated by red arrows. Scale bar: 10 μm. (B) The corresponding SRS spectra of pure cholesteryl oleate and glyceryl trioleate chemicals and CE/TG in high/low OS GBM tissues within the C-H vibrational region (2850-3000 cm⁻¹). Intensity normalized to arbitrary units (a.u.). (C) The representative maps of CE percentage in high OS and low OS IDH-wildtype GBM tissues. Color gradients (0-100%) reflect spatial CE level, with high OS IDH-wildtype GBM tissues demonstrating significantly higher CE accumulation. (D) CE percentage quantification in 89 IDH-wildtype GBM patient IDH-wildtype GBM tissues. Error bars indicate SEM (*n* > 10 samples/group). One-way ANOVA, \**P* < 0.05, \*\**P* < 0.01, and \*\*\**P* < 0.001. NO. represents the patient number.

### CE May be a Novel Biomarker in IDH-wildtype GBM Prognosis

We evaluated the clinical significance of CE as a novel prognostic biomarker in IDH-wildtype GBM. As shown in the heatmap profiling of Figure 3A, high OS patients clustered with higher CE level and MGMT methylation, while low OS cases exhibited lower CE level and MGMT unmethylation (Figure 3A). Additionally, we confirmed that MGMT methylation had a significant association with prognosis (HR = 0.331, 95% CI: 0.178-0.614, log-rank, *P* = 0.0047, Table 1 and Figure S4), consistent with prior evidence highlighting MGMT methylation as a predictor of TMZ sensitivity.^8^ Meanwhile, EGFR amplification, CDKN2A/B deletions showed no significant prognostic value (Table 1 and Figure S4), aligning with prior evidence demonstrating the lack of prognostic validity for EGFR/CDKN2A/B alterations.^40–42^ Notably, we found a strong association between CE and OS by Kaplan-Meier (KM) survival analysis (log-rank, *P* = 0.0003; Figure 3B). Specifically, patients with CE level ≥ 40% exhibited markedly improved OS compared to those with CE level < 40% (HR = 0.072, 95% CI: 0.038-0.134, log-rank, *P* < 0.0005; Figure 3C and Table 1). Collectively, these data suggested that CE may be a novel biomarker to assess IDH-wildtype GBM prognosis beside MGMT methylation.

**Figure 3.**
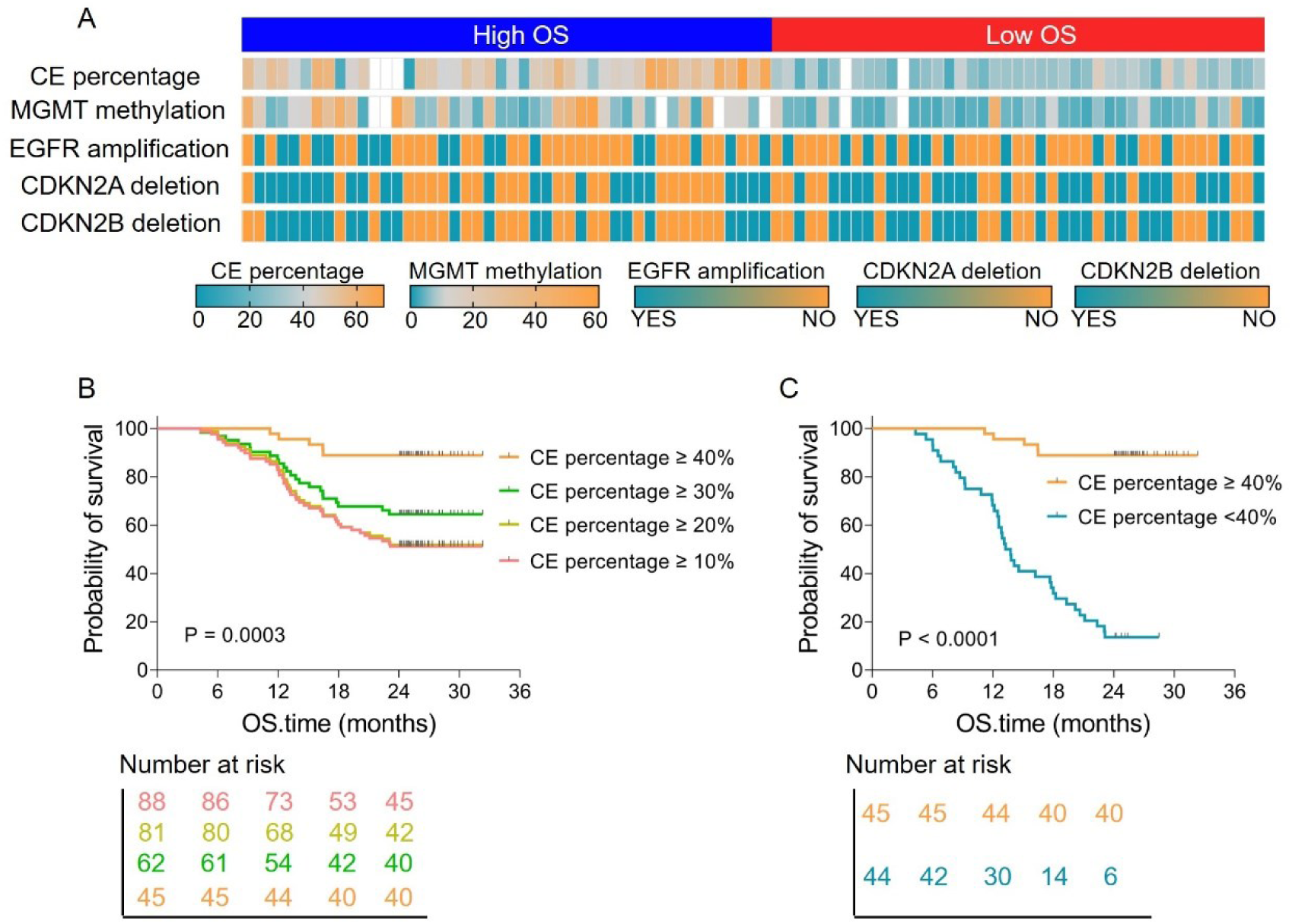
Prognostic value of CE in IDH-wildtype GBM. (A) Heatmap of CE percentage and other molecular markers (MGMT methylation, EGFR amplification, CDKN2A/B deletion) in 89 IDH-wildtype GBM patients. (B) OS analysis stratified by CE percentage thresholds (≥ 10%, ≥ 20%, ≥ 30%, ≥ 40%) in IDH-wildtype GBM. Survival probability increased progressively with higher CE levels (log-rank, *P* = 0.0003). (C) Kaplan-Meier curves comparing patients with CE ≥ 40% (*n* = 45, 5 deaths) versus CE < 40% (*n* = 44, 38 deaths). The CE ≥ 40% cohort showed markedly prolonged survival (log-rank, *P* < 0.0001), validating CE as an independent prognostic biomarker in IDH-wildtype GBM.

**Table 1.** Univariate COX regression analyses in training cohort.

| Univariate COX analysis |  |  |  |  |
| --- | --- | --- | --- | --- |
| Variable |  | Number of patients | Hazard ratio (95%CI) | P |
| OS status<sup>▽</sup> | Alive | 46 |  |  |
|  | Dead | 43 |  |  |
| CE percentage (>40% vs <40%) | > 40% | 45 | 0.072 | <0.0001**** |
|  | < 40% | 44 | (0.038-0.134) |  |
| MGMT (Methylation vs Unmethylation) | Methylation | 29 | 0.331 | 0.0047** |
|  | Unmethylation | 60 | (0.178-0.614) |  |
| EGFR amplification (YES/NO) | YES | 33 | 0.755 | 0.378 |
|  | NO | 56 | (0.412-1.3) |  |
| CDKN2A deletion (YES/NO) | YES | 50 | 1.295 | 0.406 |
|  | NO | 39 | (0.711-2.359) |  |
| CDKN2B deletion (YES/NO) | YES | 47 | 1.339 | 0.343 |
|  | NO | 42 | (0.736-2.434) |  |

▽: Two-year Overall Survival (OS)

### CE Outperforms MGMT Methylation and Prognostic Model Reveals Synergistic Prediction

We validate whether CE is a superior prognostic biomarker for IDH-wildtype GBM. To maximize the utilization of limited clinical samples and ensure balanced subgroup representation, we implemented a three-fold cross-validated design. In each iteration, approximately 58-60 patients (two folds) were allocated to the training cohort, while the remaining 28-31 patients (a single fold) served as the validation cohort. This process was repeated three times with distinct data partitions, enabling comprehensive exploration of latent law within the dataset. Based on the three-fold cross-validated design, we compared the prognostic metrics of CE and MGMT methylation. Our results suggested that CE outperformed MGMT methylation in training cohorts, with AUC (0.836 ± 0.027 vs. 0.763 ± 0.036), sensitivity (0.883 ± 0.024 vs. 0.703 ± 0.044), specificity (0.870 ± 0.001 vs. 0.781 ± 0.074), accuracy (0.864 ± 0.022 vs. 0.674 ± 0.030), negative predictive value (NPV: 0.847 ± 0.024 vs. 0.654 ± 0.074), and positive predictive value (PPV: 0.883 ± 0.048 vs. 0.695 ± 0.052) (Figure 4A and Table 2).

**Figure 4.**
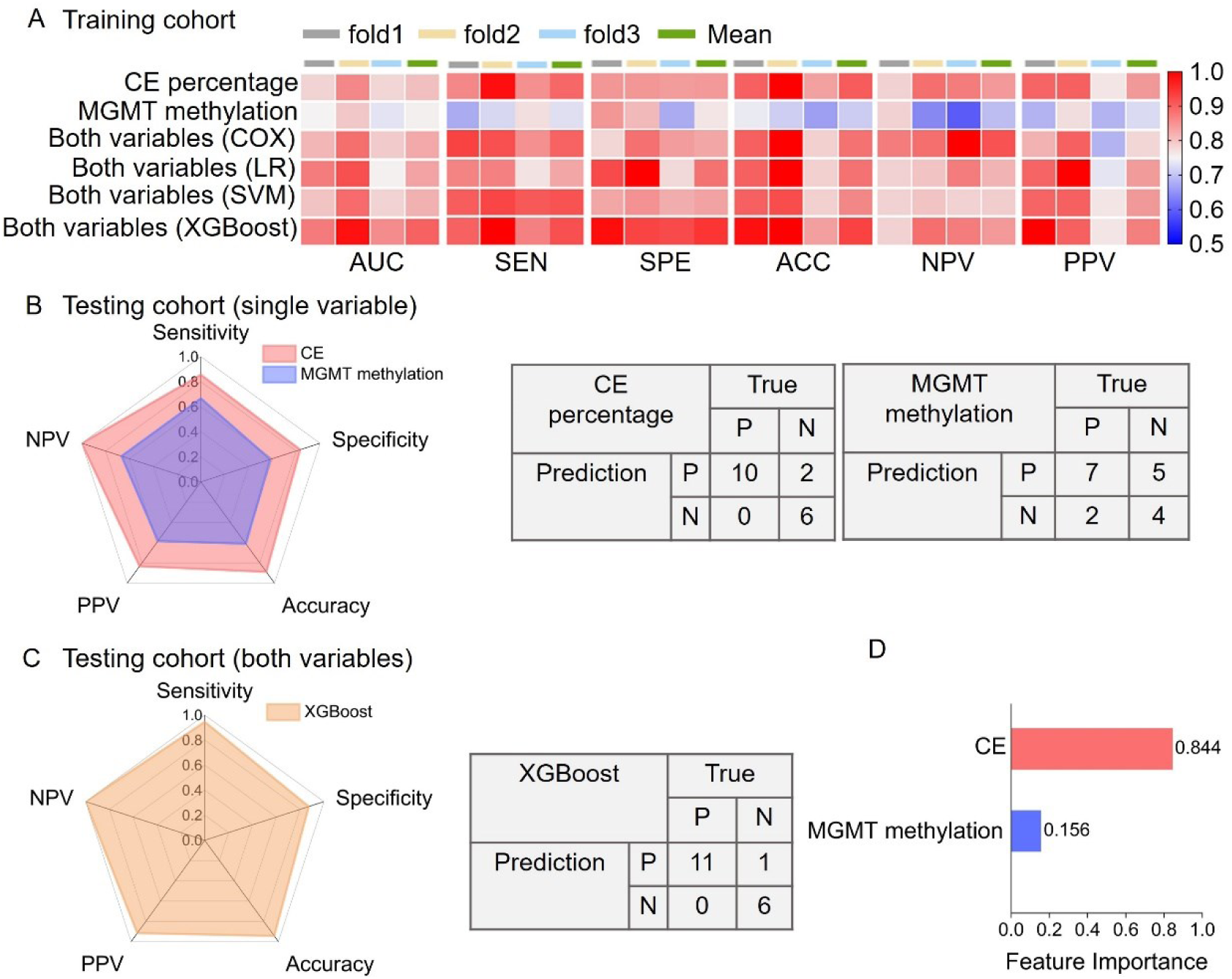
CE outperforms MGMT methylation in IDH-wildtype GBM prognosis and the integrative prognostic model. (A) Heatmap evaluating Cox regression, LR, SVM, and XGBoost models via three-fold cross-validation. Performance metrics include AUC, sensitivity (SEN), specificity (SPE), accuracy (ACC), and positive/negative predictive values (PPV/NPV). Mean represents the average of the results from the three-fold cross-validation. (B) When subsequently applying training-optimized thresholds from training to the testing cohorts, CE outperformed MGMT methylation across all metrics. (C) Validation of the XGBoost model in an independent cohort confirms maintained robust performance. (D) Feature importance analysis in the XGBoost model shows dominant contribution of CE (0.844) versus MGMT methylation (0.156).

**Table 2.**
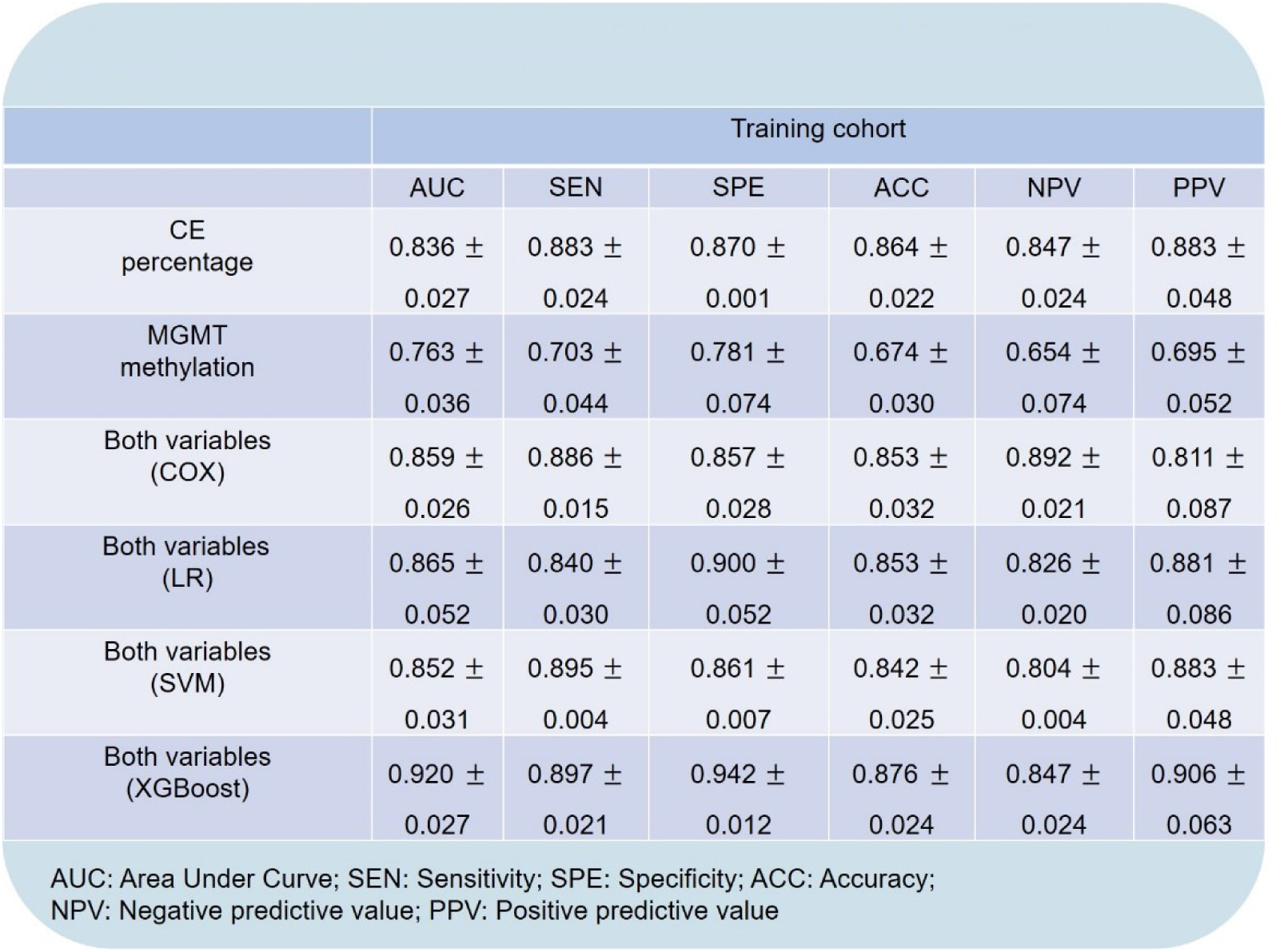
Evaluation metrics of single variable and both variables on the training Cohort.

**Table 3.**
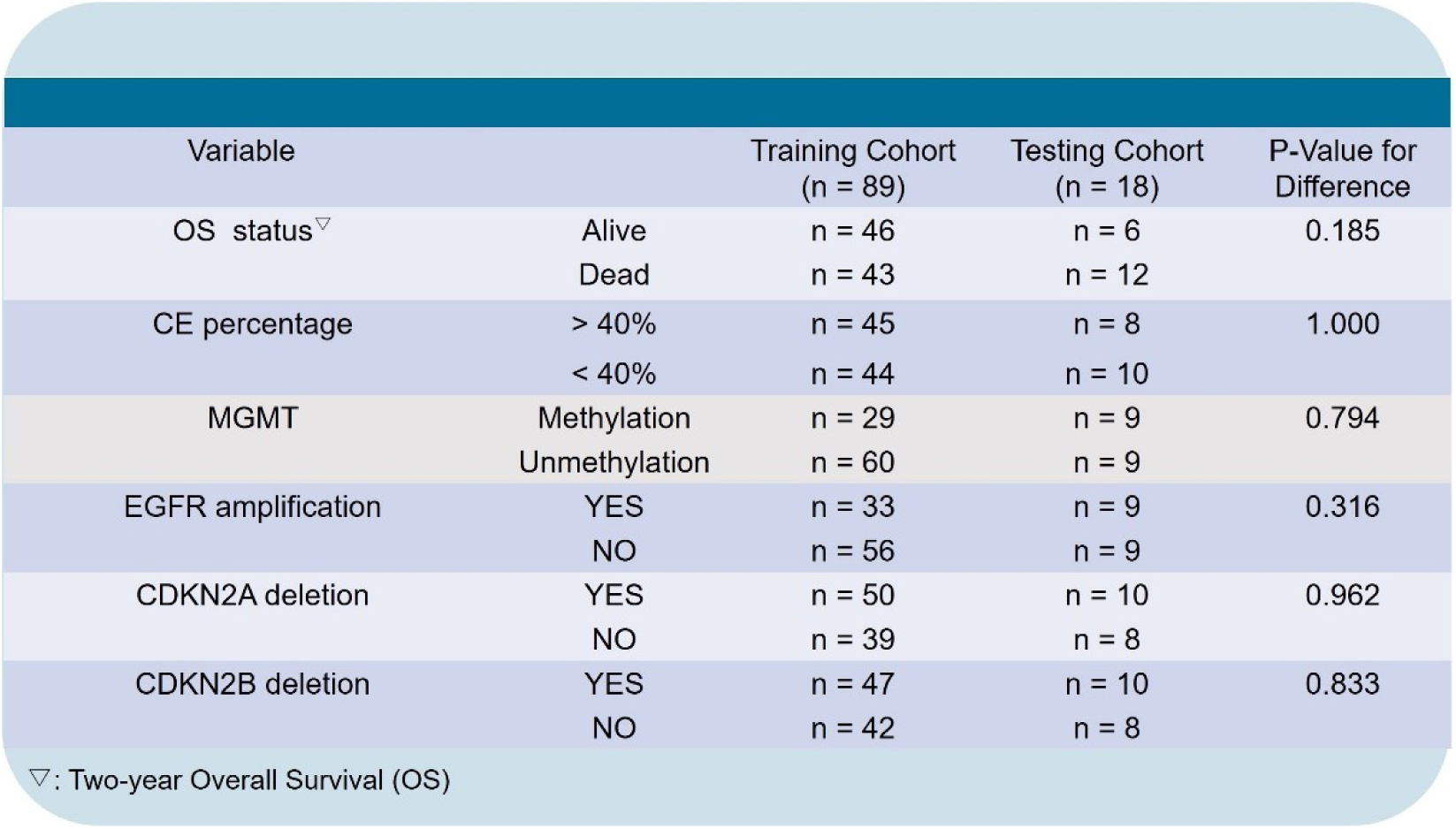
The basis data of training cohort and testing cohort.

**Table 4.** Evaluation metrics of single variable and both variables on the testing cohort.

|  | Testing cohort |  |  |  |  |
| --- | --- | --- | --- | --- | --- |
|  | SEN | SPE | ACC | NPV | PPV |
| CE | 0.856 | 0.833 | 0.889 | 1.00 | 0.833 |
| MGMT methylation | 0.667 | 0.583 | 0.611 | 0.667 | 0.583 |
| Both variables (XGBoost) | 0.946 | 0.873 | 0.944 | 1.00 | 0.917 |

AUC: Area Under Curve; SEN: Sensitivity; SPE: Specificity; ACC: Accuracy; NPV: Negative predictive value; PPV: Positive predictive value

Nonlinear interaction analysis revealed a synergistic effect between CE and MGMT methylation (Figure S5), driving the development of prognostic model that combines both biomarkers. The prognostic model was validated through three-fold cross-validation using Cox regression, logistic regression (LR), support vector machine (SVM), and extreme gradient boosting (XGBoost) models. The XGBoost model achieved exceptional performance in validation cohort, with AUC (0.920 ± 0.027), sensitivity (0.897 ± 0.021), specificity (0.942 ± 0.012), accuracy (0.876 ± 0.024), NPV (0.847 ± 0.024), and PPV (0.906 ± 0.063) (Figure 4A and Table 2). This represents a substantial improvement over single-variable model. The cyclic training-validation protocol not only minimized overfitting risks but also enhanced the generalizability of our survival prediction model, ensuring feasible clinical applicability for patient stratification and prognosis assessment.

To evaluate the generalizability, we further tested the prognostic value of the variables (CE alone, MGMT methylation alone, or a combination of both) in an independent cohort of 18 patients. There were no statistically significant differences in OS status, CE percentage, MGMT methylation, EGFR or CDKN2A/B between the training and testing cohorts (Table 3). In the testing cohort, CE maintained superior performance over MGMT methylation, which achieved higher sensitivity (0.856 vs. 0.667), specificity (0.833 vs. 0.583), accuracy (0.889 vs. 0.611), NPV (1.00 vs. 0.667), and PPV (0.833 vs. 0.583) (Figure 4B and Table 4). Moreover, the XGBoost model retained robust prognostic performance, achieving high sensitivity of 0.946, specificity of 0.873, accuracy of 0.944, NPV of 1.00, and PPV of 0.917 (Figure 4C and Table 4). Feature importance analysis revealed CE as the dominant contributor compared to MGMT methylation (Figure 4D). This study establishes CE as a clinically reliable prognostic biomarker that outperforms MGMT methylation, while their integration through machine learning uncovers unprecedented predictive accuracy.

### Gene Expression Profiling Underlying Impact of CE in IDH-wild GBM Prognosis

To investigate the regulatory mechanism of cholesterol metabolism and poor prognosis, we conducted a systematic analysis using clinical and transcriptomic data from 160 IDH-wild GBM patients in the TCGA database. The volcano plots displayed the differentially expressed genes (DEGs) in the HR <1 and HR >1 (Figure S6), and the common DEGs were then used for further functional analysis based on gene ontology (GO) (Figure S7). Subsequently, we focused on investigating genes related to cholesterol metabolism. Among them, high expression of genes involved in cholesterol efflux (ABCA5/PLTP/PPARG), hydrolysis (CYP27A1), transport (NPC2) and inhibition of de novo synthesis (PPARG) was strongly associated with shorter OS by KM survival analysis (Figures 5A-5F and S8).

**Figure 5.**
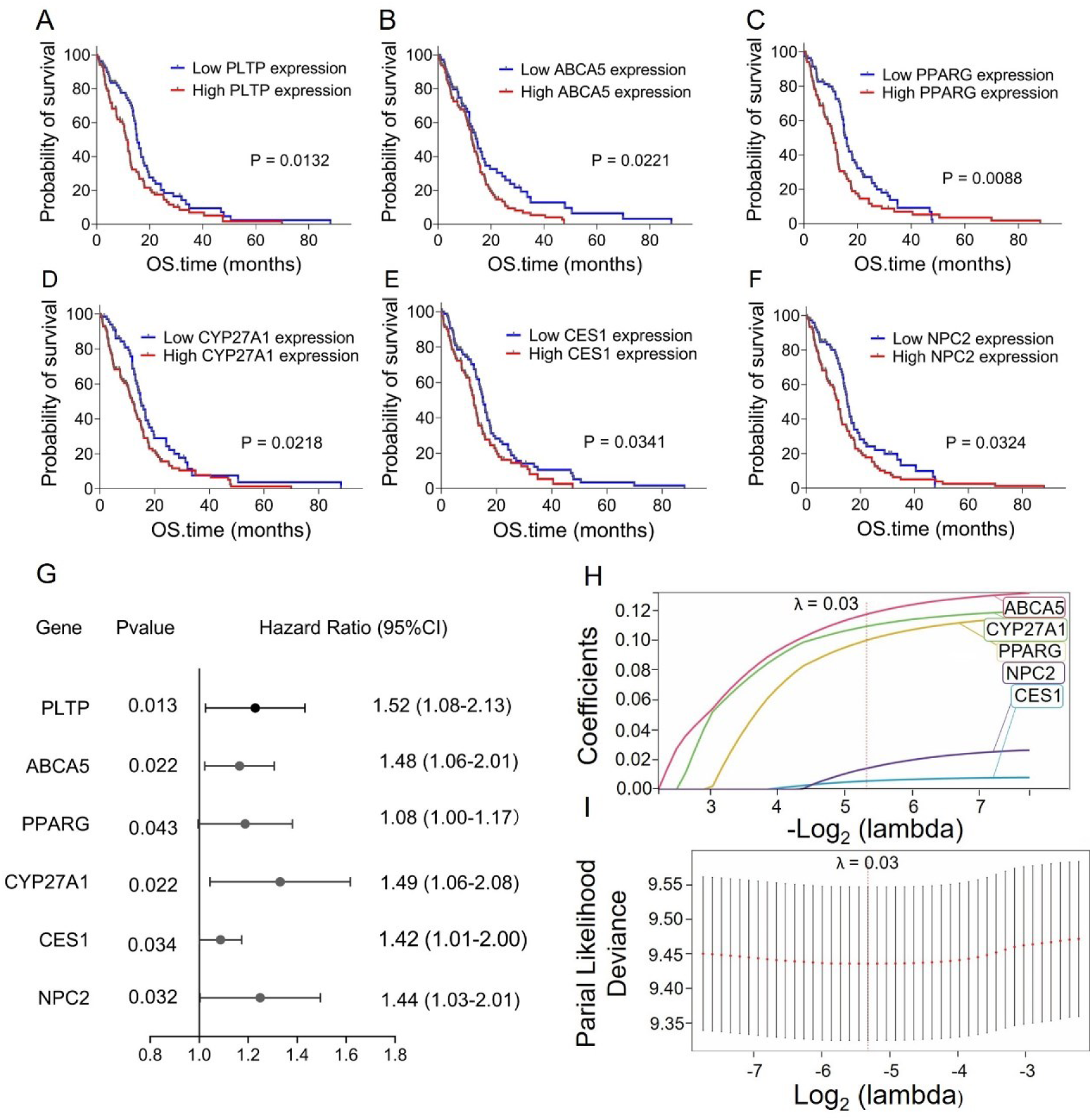
Validation of cholesterol metabolism-related gene risk signature in IDH-wildtype GBM prognosis. (A-F) Kaplan-Meier curves of OS stratified by low/high expression of cholesterol metabolism-related genes (PLTP, ABCA5, PPARG, CYP27A1, CES1, NPC2) in 160 IDH-wildtype GBM patients from TCGA. Log-rank test: PLTP (*P* = 0.013), ABCA5 (*P* = 0.022), CYP27A1 (*P* = 0.022), NPC2 (*P* = 0.032), CES1 (*P* = 0.034) and PPARG (*P* = 0.043). (G) Univariate Cox analysis evaluates the prognostic value of the cholesterol metabolism-related genes in terms of OS. (H-I) Lasso Cox regression selected four genes (ABCA5, PPARG, CYP27A1, NPC2) with non-zero coefficients.

Since single gene only reflects unidimensional tumor biology, we established a cholesterol metabolism-guided risk score model to evaluate the synergistic prognostic effect of key genes. To prevent overfitting and enhance the risk score model’s generalization ability, we further identified four genes (ABCA5, PPARG, CYP27A1, NPC2) contributing most significantly to prognosis using the LASSO regression analysis (Figures 5G-5I). Based on the risk score model integrating survival status and survival time, patients were stratified into high-risk and low-risk group using the median risk score (Figure 6A). KM analysis confirmed the model’s prognostic effect for OS (log-rank, *P* < 0.005; Figure 6B), which was further validated by the GSE74187 cohort (log-rank, *P* < 0.05, Figure 6C). Notably, high-risk patients showed marked upregulation of genes related to cholesterol transport (NPC2), efflux (ABCA5/PPARG), hydrolysis (CYP27A1) and inhibition of de novo synthesis (PPARG) (Figures 6A and S9). Furthermore, given the pivotal biological role of cholesterol metabolism in IDH-wildtype GBM prognosis, the connection between risk score and cholesterol metabolism-related genes was thoroughly studied. The results illustrated that patients with an elevated risk score showed a positive correlation with ABCA5 (*r* = 0.51), PPARG (*r* = 0.43), CYP27A1 (*r* = 0.59), and NPC2 (*r* = 0.47) (Figures 6D-6G).

**Figure 6.**
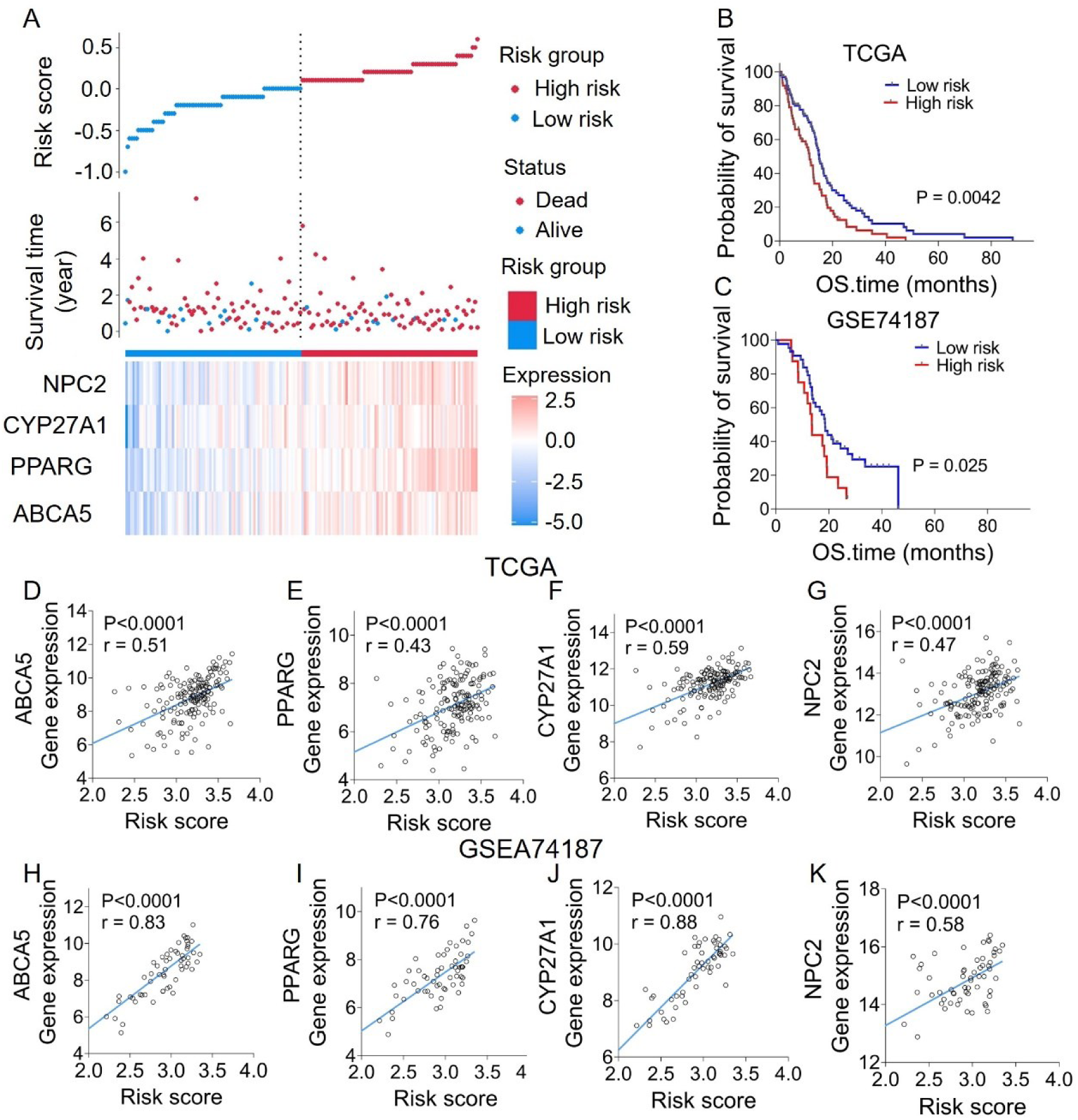
Construction and validation of the risk score model for IDH-wildtype GBM. (A) Risk score distribution (top, gradient bar), survival status (middle, red: dead; blue: alive), and heatmap of four prognostic gene signatures (ABCA5, PPARG, CYP27A1, NPC2) in 160 IDH-wildtype GBM patients from TCGA. Higher risk scores correlate with reduced survival time (OS.time, year). (B-C) Kaplan-Meier survival curves stratified by low-risk (blue) and high-risk (red) groups in TCGA (B, log-rank, *P* = 0.0042) and GSE74187 (C, log-rank, *P* = 0.025) cohorts. (D-G) Scatterplots showing positive correlations between risk score and expression of ABCA5 (*r* = 0.51, *P* < 0.0001), PPARG (*r* = 0.43, *P* < 0.0001), CYP27A1 (*r* = 0.59, *P* < 0.0001), and NPC2 (*r* = 0.47, *P* < 0.0001) in TCGA. (H-K) Validation of gene-risk score correlations in GSE74187: ABCA5 (*r* = 0.83, *P* < 0.0001), PPARG (*r* = 0.76, *P* < 0.0001), CYP27A1 (*r* = 0.88, *P* < 0.0001), NPC2 (*r* = 0.58, *P* < 0.0001).

These results were further validated by GSE74187 cohort (Figures 6H-6K). These findings together indicate that cholesterol metabolic reprogramming promotes tumor malignancy by activating cholesterol efflux and hydrolysis pathways and inhibiting de novo synthesis, positioning cholesterol-related gene signatures as potential clinical biomarkers for IDH-wildtype GBM risk stratification.

## DISCUSSION

Aberrant cholesterol metabolism is recognized as a hallmark of GBM, however, the prognostic role of CE remains undefined. Through high spatial resolution SRS imaging, we revealed that CE not only serves as an independent prognostic biomarker but also outperforms conventional MGMT methylation in prognostic prediction for IDH-wildtype GBM. Furthermore, the prognostic model combining CE and MGMT methylation uncovered synergistic prognostic effects. Mechanistic analyses further revealed that upregulation of genes involved in cholesterol transport, efflux, hydrolysis, and inhibition of de novo synthesis contributed to reduced CE in poor prognosis GBM. As discussed below, these findings improve current understanding of the role of CE in IDH-wildtype GBM prognosis.

First, our study demonstrates the clinical value of CE as a novel biomarker for prognostic evaluation in IDH-wildtype GBM. While MGMT methylation guides current clinical decisions, its limitations are increasingly apparent. There is high variability in testing methods, as each assay evaluates methylation status differently, making an appropriate cut-off difficult to define.^14, 43, 44^ Furthermore, these approaches exhibit suboptimal diagnostic accuracy, as evidenced by AUC values frequently below 0.8, alongside compromised sensitivity and specificity.^16–18^ In contrast, CE outperforms MGMT methylation, characterized by superior AUC (0.832). The validation of CE as a prognostic biomarker deserves further confirmation through a large, multi-institutional cohort.

Take advantage of synergistic effects between CE and MGMT methylation, we developed a prognostic model. The prognostic model (AUC = 0.920) revealed complementary roles of CE and MGMT methylation: CE may remodel tumor plasticity via lipid metabolism, while MGMT methylation enhances alkylating agent sensitivity through DNA repair defects. This cross-pathway synergy resembles imaging-molecular integration strategies like relative cerebral blood volume (rCBV) combined with MGMT model, yet our model elevates AUC to 0.920 by incorporating metabolic dimensions, markedly outperforming conventional combinatorial models (AUC = 0.85).^45^ This study provides an innovative and clinically feasible prognostic model for GBM prognosis.

Second, this study uncovers a potential mechanistic link between cholesterol metabolism and IDH-wildtype GBM prognosis. Previous studies have reported that CE is prevalent in GBM and can be used as potential targets, characterized by disrupted uptake, and efflux dynamics, leading to abnormal CE depletion and subsequent suppression of tumor proliferation.^23–25^ These studies included patients with IDH-mutant tumors historically classified as GBM, which are now redefined as astrocytoma, IDH-mutant, WHO 4.^3,4^ Importantly, emerging evidence demonstrates a correlation between CE level and IDH status, with lower CE observed in more aggressive IDH-wild type GBM.^26,46,47^ Notably, our study revealed that such remarkable CE reduction only occurred in the low OS IDH-wildtype GBM. We further found that differential

expression of cholesterol metabolism-related genes was strongly associated with patient outcomes. In high-risk patients with ABCA5 overexpression, excessive cholesterol efflux may destabilize lipid raft structures, impairing membrane signaling (e.g., EGFR/PI3K pathways) and accelerating tumor progression.^48, 49^ PPARG inhibits de novo cholesterol synthesis (e.g., via HMGCR degradation) and promotes efflux, but may activate pro-inflammatory cytokines that exacerbated the immunosuppressive tumor microenvironment (TME).^50,51^ NPC2 mediates free cholesterol transport from endosome/lysosome complexes to the endoplasmic reticulum (ER) and plasma membrane.^52,53^ CYP27A1 catalyzes cholesterol conversion to bile acids, while its oxysterol product 27-hydroxycholesterol (27-HC) activates the LXR pathway, upregulating cholesterol efflux and downregulating LDLR expression to reduce cholesterol levels.^54,55^ These pathways collectively account for the reduction in CE observed in IDH-wildtype GBM with poor prognosis, mediated through enhanced efflux (ABCA5) and hydrolysis (PPARG/CYP27A1). Although the biological mechanisms of cholesterol metabolism require further elucidated, these findings still uncover possible mechanistic insights into the link between cholesterol metabolism and IDH-wildtype GBM aggressiveness.

Third, SRS imaging, with its label-free, non-destructive, and high-resolution capabilities, demonstrates significant potential for clinical translation in oncology. Traditional genomic and proteomic analyses for GBM involves in RNA extraction from biopsy samples, polymerase chain reaction (PCR) amplification, and microarray-based expression profiling, which make the whole process very lengthy and prone to structural damage during sample processing.^56–58^ In contrast, SRS enables direct submicron-resolution imaging of unprocessed biological tissues without staining or complex preprocessing, eliminating the need to transfer specimens out of the operating room to a pathology laboratory for sectioning, mounting, dyeing, and interpretation. Moreover, because tissue preparation for SRS microscopy is minimal, key tissue architectural details commonly lost in smear preparations and cytologic features often obscured in frozen sections are preserved. Meanwhile, SRS could achieve standardized diagnostic workflows through quantitative analysis.^59^ This capability is particularly advantageous for rapid intraoperative pathological assessment, such as real-time delineation of tumor boundaries during brain tumor resection, aiding surgeons in optimizing resection margins and minimizing neurological damage.^60^ Although we currently used large-scale equipment, advancements in fiber laser technology have enabled device miniaturization and operational simplification. For instance, deep learning-integrated SRS facilitates rapid intraoperative diagnosis of brain tumors and guides biopsy procedures.^61,62^ By standardizing data acquisition and quantitative analysis, SRS is poised to become a cornerstone tool for GBM molecular subtyping, ultimately improving patient outcomes and reducing healthcare costs.

In summary, with the superior capability of quantitative chemical imaging, our SRS spectral imaging revealed CE correlation with prognosis in IDH-wildtype GBM, which have not been documented before. Meanwhile, we provide a 41.08% CE level that can serve as a discriminator to predict IDH-wildtype GBM prognosis. When combined with the clinically used MGMT methylation, CE offers better predictive outcomes than MGMT methylation alone. Furthermore, this work unravels the hidden relationship between prognosis and cholesterol metabolism in IDH-wildtype GBM, which may offer new opportunities for the diagnosis and treatment of poor-prognosis GBM.

## MATERIALS AND METHODS

### Human GBM Tissue Specimens

This study includes 107 patients diagnosed with IDH-wildtype GBM. All tissues specimens were obtained from the patients undergone resection surgery in Neurosurgery Department of Beijing Tiantan Hospital. The tissues specimens were snap-frozen in liquid nitrogen within 10 min of surgery excision. Each sample was ∼100 mg. Frozen tissue samples were embedded in Optimal Cutting Temperature media for further sectioning into pairs of neighboring slices, with one unstained 20-µm slice for spectroscopic imaging and the other neighboring 7-µm slice for H&E staining. Pathological examination was performed by experienced pathologists. The rest tissue samples were stored in −80 ℃ refrigerator. This study was approved by the institutional review board.

### Chemical Reagents

Pure cholesteryl oleate (Sigma Grade, ≥ 99%) and glyceryl trioleate (Sigma Grade, ≥ 99%) were purchased from Sigma-Aldrich (St. Louis, MO).

### Label-free SRS Imaging

Label-free SRS imaging was performed on the Multimodal Nonlinear Optical Microscopy System (UltraView, Beijing Wellmed Medical Diagnostics & Laboratory Co., Ltd), with a dual-output femtosecond (fs) laser source (Insight X3, Spectra-Physics) providing two synchronized beams with a repetition rate of 80 MHz was employed for imaging. The pump beam was tunable from 680 to 1300 nm with a pulse width of 120 fs, and the Stokes beam was fixed at 1045 nm with a pulse width of 220 fs. An acoustic-optic (AOM, Z-1064-100-L-D0.5-T-A-K, CSRayzer Optical Technology Co., Ltd.) was used to modulate the Stokes beam at a frequency of 2.7 MHz. After combination, two collinear beams were coupled to dichroic mirror (DMLP1000, Thorlabs) and delivered to a commercial upright laser scanning microscope. Then, a 60X Water objective (NA = 1.2, UPLSAPO, Olympus) was used to focus two beams on the sample, and an oil immersion condenser (NA = 1.4, U-AAC, Olympus) was used to collect the signals from the sample. Two filters (ET980SP, ET795/150, Chroma) were used to filter out the Stokes beam, the pump beam was detected by a photodiode (S3994-01, Hamamatsu), and the SRS signal was extracted by a lock-in amplifier (MFL1, Zurich Instrument).

To detect the vibrational bands of CH_2_ (2850 cm^−1^) and the characteristic band for cholesterol (2870 cm^-1^), the wavelength of the pump beam was tuned to 801 and 804 nm, respectively. The power of the pump beam at the specimen was maintained at ∼13 mW, and the power of the Stokes beam was kept at ∼ 60 mW. SRS images containing 400 × 400 pixels were collected with a pixel dwell time of 30 µs, and then analyzed using ImageJ and custom-written MATLAB scripts.

Spontaneous Raman spectra were acquired with a Raman micro-spectrometer (DR316B-LDC-DD, Andor), which was mounted to the side of the microscope. The pump beam from a picosecond laser (Applied Physics & Electronics, picoEmerald^TM^ S) was tuned to ∼707 nm and coupled into the microscope for Raman spectra measurements. The excitation power at the sample was ∼32 and ∼60 mW for pure chemicals and tissue specimens, respectively. Each Raman spectrum ranging from 600 to 3050 cm^−1^ was acquired in 20 s for pure chemicals and 60 s for tissue specimens.

### Quantification of CE Percentage

We used the ratio metric approach based on the height ratio of the 2870 cm^-1^ peak to the 2850 cm^-1^ peak (I_2870_/I_2850_) to quantify the CE percentage according to the literature ^38, 39^. The measurements of the TG/CE emulsions and establishment of calibration curve were shown in Figure S2. The CE percentage was linearly proportional with the Raman height ratio of 2850 cm^-1^ and the characteristic band for cholesterol 2870 cm^-1^ (I_2870_/I_2850_). Specifically, for this system, I_2870_/I_2850_ = 0.0031 × CE percentage (%) + 0.8162. The quantification procedure was illustrated in Figure S3. CE percentage for each tissue specimen was obtained by averaging 6-8 images.

### XGBoost Model

XGBoost is typically described as an efficient gradient boosting algorithm that enhances the performance of predictive models by constructing a series of decision trees. Implemented using the XGBoost package with hyperparameter optimization via grid search. Key parameters included a maximum tree depth of 50, learning rate (*η*) of 0.01, and estimator of 300 to prevent overfitting. Gradient-boosted trees were trained to minimize the logistic loss function, prioritizing feature importance rankings for CE and MGMT methylation. Thus, we performed three-fold cross validation in training cohort and evaluated the true generalizability of the model in testing cohort.

### Cox Regression Model

The Cox regression model is a semi-parametric regression model proposed by British statistician D.R. Cox in 1972. Fitted via the lifelines package with Breslow’s method for handling tied events. Proportional hazards assumptions were validated using Schoenfeld residuals. Thus, we performed 3-fold cross validation design.

### LR Model

LR is used to predict the probability of a binary outcome, with a simple model that is easy to understand and interpret; it can handle both categorical and continuous variables; and it outputs probability values, facilitating decision-making. Built using scikit-learn with L1 regularization (*C* = 0.5). Thus, we performed 3-fold cross validation design.

### SVM Model

SVM is supervised machine learning models that try to separate binary datasets using a simple linear function. Utilized a radial basis function (RBF) kernel with penalty parameter *C* = 5 and kernel = rbf. Thus, we performed three-fold cross validation in training cohort.

### Three-Fold Cross Validation

The study cohort comprised 89 patients (46 alive and 43 dead; alive-to-dead ratio = 1.07). To maintain the alive-to-dead balance across folds, the cohort was randomly stratified into three equal-sized folds. Each fold comprised 29-30 patients with a preserved alive-to-dead ratio of 1.07 (alive 15-16 vs. dead 14-15 per fold). In each iteration, approximately 58-60 patients (two folds) were allocated to the training cohorts, with the remaining 29-31 patients (a single fold) serving as the validation cohorts. This process was repeated three times with distinct data partitions, and performance metrics were averaged across all validation cycles.

### Single Biomarker Analysis by Three-Fold Cross Validation

For individual biomarkers (CE or MGMT methylation), the feature values in the training cohort were directly utilized as continuous predictors. The optimal classification cutoff was determined by identifying the point on the ROC curve that maximized the difference between the true positive rate (TPR) and false positive rate (FPR). This cutoff was subsequently applied to the validation cohort to predictive outcomes. Notably, AUC was computed exclusively on the training set due to the absence of continuous predictive values in the validation set (only predictions were generated from the fixed cutoff), precluding ROC curve construction for the validation cohort. All other metrics (sensitivity, specificity, etc.) were evaluated on the validation set. Using the training cohort-optimized threshold, we tested the single variable on an independent cohort of 18 patients.

### The Prognostic Model Analysis by Three-Fold Cross Validation

For the prognostic model combining CE and MGMT methylation, nonlinear regression models were performed using both biomarkers as independent variables and survival status (coded as 1 for alive and 2 for dead) as the dependent variable. After deriving model parameters from training data, predicted values were generated for validation cohorts. ROC curves were subsequently constructed using these continuous predictions, enabling AUC calculations and optimal threshold determination equivalent to the single-marker methodology. All validation outcomes were aggregated across three independent data splits for final performance reporting.

### Feature Importance Assessment

To evaluate the contribution of input features to predictive performance, we computed feature importance scores using XGBoost’s built-in metric. Feature importance was quantified via the “gain” method, which calculates the average improvement in predictive accuracy (reduction in loss) generated by each feature across all decision trees in the ensemble. This approach assigns higher importance to features that most significantly enhance node purity during splits. The two factors assessed CE and MGMT methylation-were ranked according to their normalized importance scores (summing to 1), providing a relative measure of their predictive influence. All importance values were derived from the model’s native “get_score” function with importance-type = gain.

### Clinical Performance Metrics

Evaluation metrics included AUC, sensitivity, specificity, accuracy, PPV, and NPV. In a binary classification problem (as those we consider are), Death is considered to be the ‘positive’ and survival is considered to be the ‘negative’ class. True positive (TP) presented patients who died (ground truth) and were correctly predicted as high-risk (death) by the model. The false positive (FP) correspond to patients who survived (ground truth) but were incorrectly classified as high-risk (death). The false negative (FN) correspond to patients who died (ground truth) but were erroneously classified as low-risk (survival). The true negative (TN) correspond to patients who survived (ground truth) and were correctly predicted as low-risk (survival). PPV = TP/(TP+FP)-Probability that a predicted high-risk patient will die. NPV = TN/(TN+FN) - Probability that a predicted low-risk patient will survive.

### Confusion Matrices Definition

Confusion matrices are useful for describing the predictive capabilities of machine learning models. The rows in a confusion matrix correspond to the classes in the dataset, while the columns correspond to the predicted classes. In the matrix, the diagonal is made up of all true positive predictions the model has produced. In a binary classification problem (as those we consider are), Death is considered to be the ‘positive’ and survival is considered to be the ‘negative’ class. TP is listed in the top left corner of the confusion matrix. The FP is listed in the top right corner of the confusion matrix. The FN is listed in the bottom left corner of the confusion matrix. The TN is listed in the bottom right corner of the confusion matrix. In our 3-fold cross-validation, we computed distinct confusion matrices for each model on their respective validation sets and summed them up to measure the overall model performance across the entire dataset.

### Public Database

TCGA, a project supported by the National Cancer Institute (NCI) and National Human Genome Research Institute (NHGRI), has generated comprehensive, multi-dimensional maps of the key genomic changes in various types of cancers. The GBM database in TCGA-GBM, including gene expression and survival data, was obtained from the UCSC Xena browser (https ://xenabrowser.net). We obtained 160 IDH-wildtype GBM cases, who contained survival status, gene expression and survival data.

GSEA was conducted to assess whether there were considerable variations in the set of genes expressed between the high-risk and low-risk cohorts. The GBM database in GSEA74187, including gene expression and survival data, was obtained from the from the Gene Expression Omnibus (GEO) database (http://www.ncbi.nlm.nih.gov/geo/). We obtained 60 GBM cases, who contained survival, gene expression and survival data.

### Differentially Expressed Genes (DEGs) Analysis

The principle of differential analysis was to first convert the count matrix into an object, then each gene gets assigned the same dispersion estimate, then performs pair-wise tests for differential expression between two groups, and finally takes the output using the False Discovery Rate (FDR) correction, and returns the top differentially expressed genes. The parameters set for differential expression analysis were FDR < 0.05 with |Log2FC| > 2. Subsequently, Gene Ontology (GO) analysis was used to do the biological function analysis. R programming language was used to analyze the correlation between survival data and other genes expression with the Pearson correlation analysis. The figures were plotted with the GraphPad prism9.

### Statistical Analysis

All data were shown as mean ± standard error of the mean (SEM). One-way ANOVA was used for comparisons among groups. Survival analyses used log-rank tests (KM curves). *P* < 0.05 was considered statistically significant.

## Supporting information

Supplemental Table 1, and will be used for the link to the file on the preprint site.

## Acknowledgments

This work was supported by National Natural Science Foundation of China (81930048; 62027824) and The Capital Health Research and Development of Special (2022-2-2047).

## Author contributions

All authors made substantial contributions to the conception or design of the study; the acquisition, analysis, or interpretation of data; or drafting and revising the manuscript. All authors approved the manuscript. All authors agree to be personally accountable for individual contributions and to ensure that questions related to the accuracy or integrity of any part of the work are appropriately investigated, resolved, and the resolution documented in the literature. N.N.W. conducted the SRS image acquisition, data analysis and manuscript preparation. J.J.W conducted tissue preparation and participated in SRS imaging acquisition. J.L.L was responsible for writing the model code. J.Q.Z. was responsible for SRS image acquisition. B.Y. participated in SRS imaging acquisition. P.W., S.H.Y., and N.J. were responsible for experimental design. N.W. wrote the manuscript. S.H.Y. and N.J. wrote and edited the manuscript. All authors contributed to the study and critically reviewed the manuscript.

## Data and materials availability

All data are available in the main text or the supplementary materials.

