## Supplemental Table 1, and will be used for the link to the file on the preprint site. for "Cholesteryl Ester as a Prognostic Biomarker In IDH-wildtype Glioblastoma"

[‡]Equal contribution.

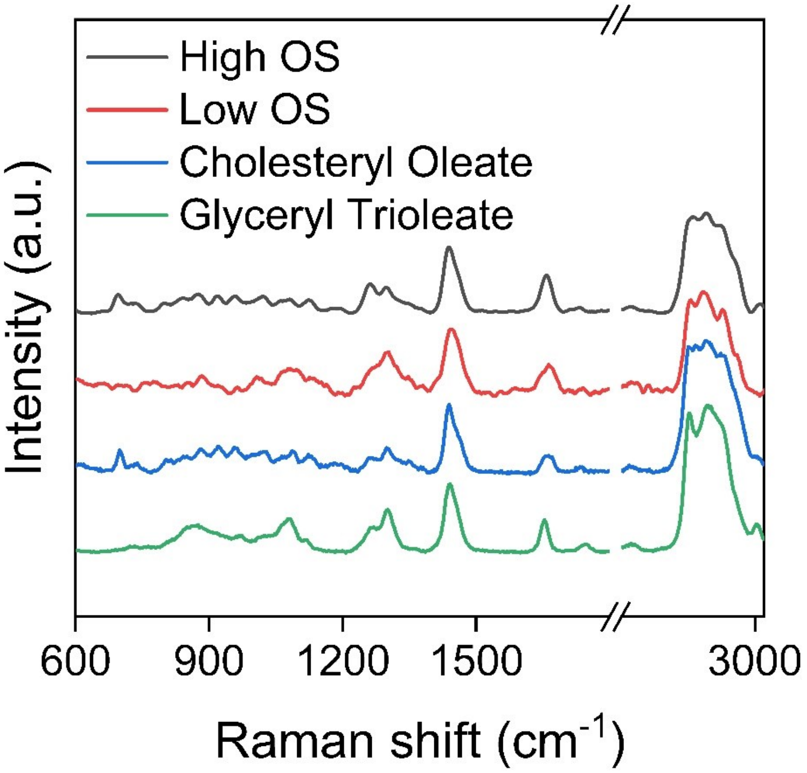


**Supplementary Figure 1. Spontaneous Raman of pure cholesteryl oleate (blue) and glyceryl trioleate (green) chemicals, and CE/TG extracted from high OS (black) and low OS (red) IDH-wildtype GBM tissues.**

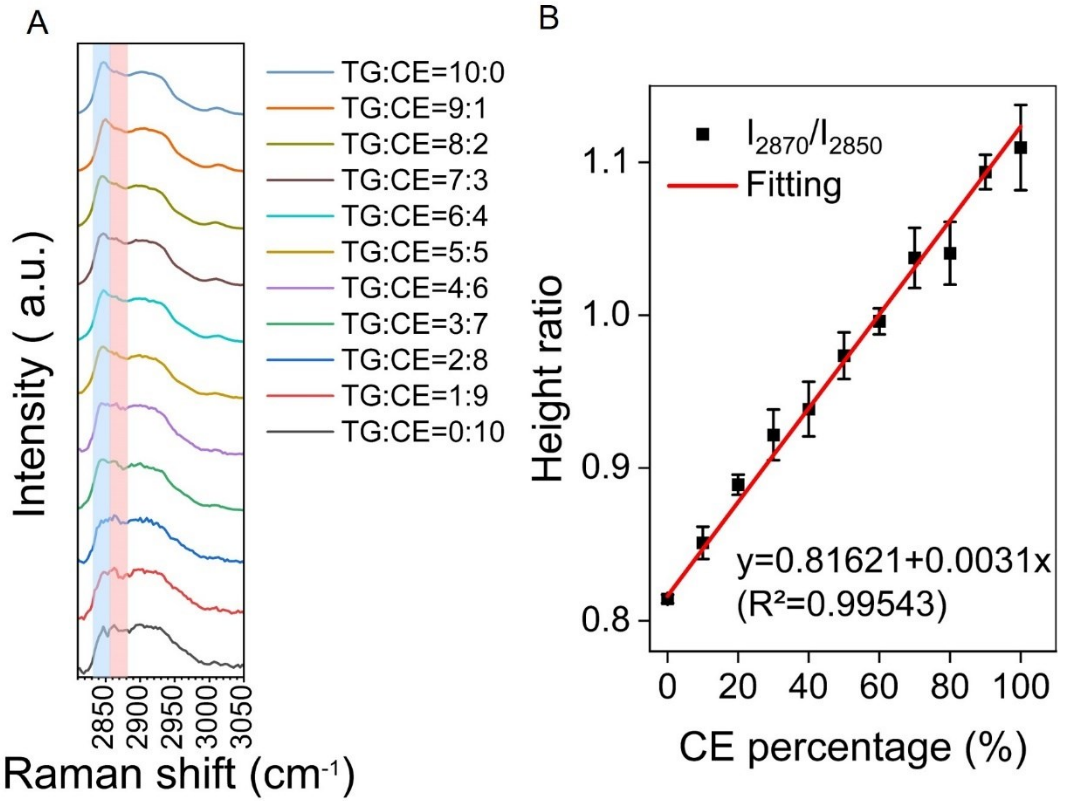


**Supplementary Figure 2**. **Quantitative calibration of CE percentage in neutral lipid mixtures using Raman spectroscopy.** (A) Raman spectra of CE and TG emulsion standards with TG : CE molar ratios ranging from 10:0 to 0:10 (color-coded). Spectra were normalized to the C-H stretching peak at 2850 $cm^{-1}$ (dashed vertical line). Distinctive spectral shifts at 2870 $cm^{-1}$ (CE-enriched phase) and 2850 $cm^{-1}$ (TG-enriched phase) were resolved. (B) Linear calibration curve (red line) derived from the 2870/2850 $cm^{-1}$ peak height ratio (black squares ± SEM). The fitted equation $y = 0.81621 + 0.0031x$ ($R^2 = 0.99543$) enabled precise CE percentage quantification (0-100% range). This validated model was subsequently applied to CE quantification in GBM tissue samples.

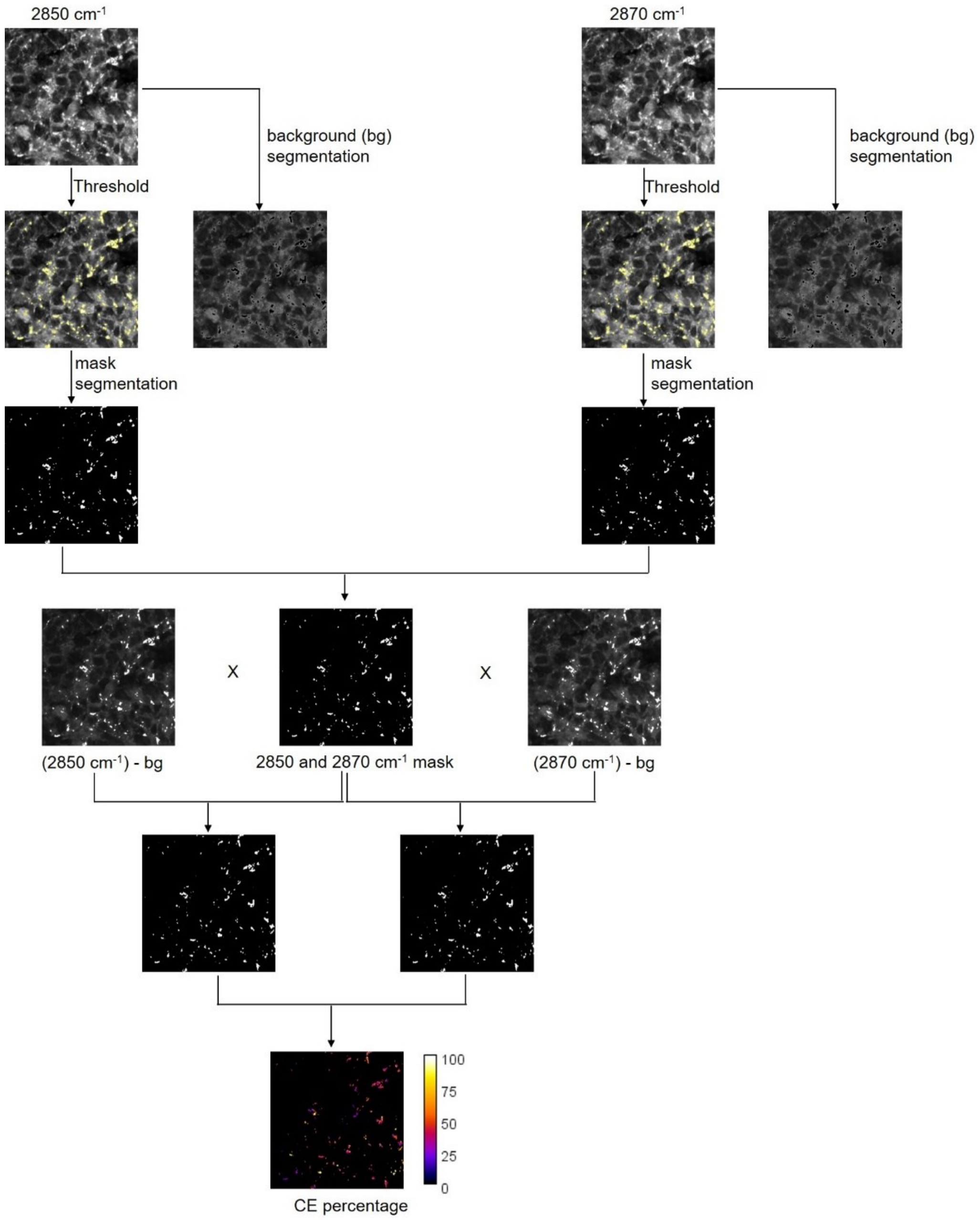


**Supplementary Figure 3. The workflow of image processing and the quantification of CE percentage, in representative SRS images of human IDH-wildtype GBM tissue**. CE in the cell can be selected based on their significantly higher signal intensities compared to other cellular compartments. By using ImageJ 'Threshold' function, raw images were converted to binary images, which were used for mask segmentation. The images of 2850 cm$^{-1}$ and 2870 cm$^{-1}$can be obtained by multiplying the mask image with the SRS image of 2850 cm$^{-1}$ and 2870 cm$^{-1}$, respectively. Quantification of CE percentage in tissues was then calculated based on the established calibration curve shown in Fig. S2B.

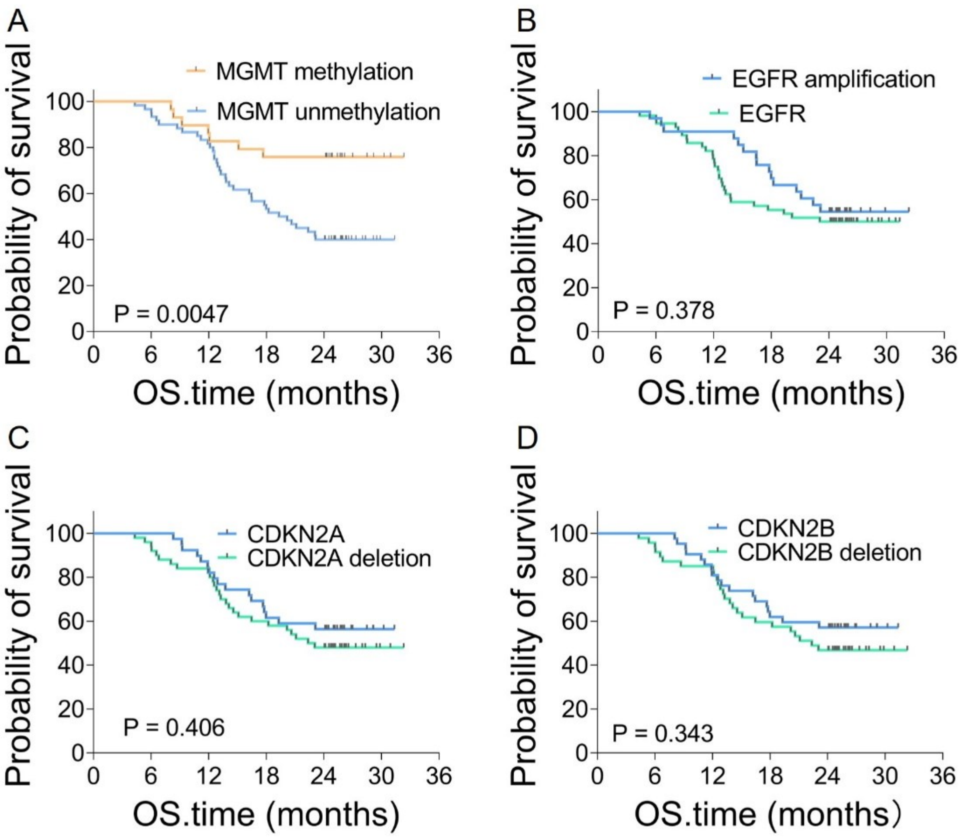


**Supplementary Figure 4. Kaplan-Meier survival analysis of molecular markers in IDH-wildtype GBM.** (A-D) Survival curves stratified by molecular status in 89 IDH-wildtype GBM patients: (A) MGMT methylation (methylated: $n$ = 29 vs. unmethylated: $n$ = 60, log-rank, $P$ = 0.0047); (B) EGFR amplification (Yes: $n$ = 33 vs. No: $n$ = 56, P = 0.378); (C) CDKN2A deletion (Yes: $n$ = 50 vs. No: $n$ = 39, $P$ = 0.406); (D) CDKN2B deletion (Yes: $n$ = 47 vs. No: $n$ = 42, $P$ = 0.343). MGMT methylation GBM patients exhibited significantly prolonged OS, while EGFR, CDKN2A, and CDKN2B status showed no statistically significant prognostic association (log-rank test, df = 1). Axes: $x$ = survival time (months); $y$ = survival probability (%).

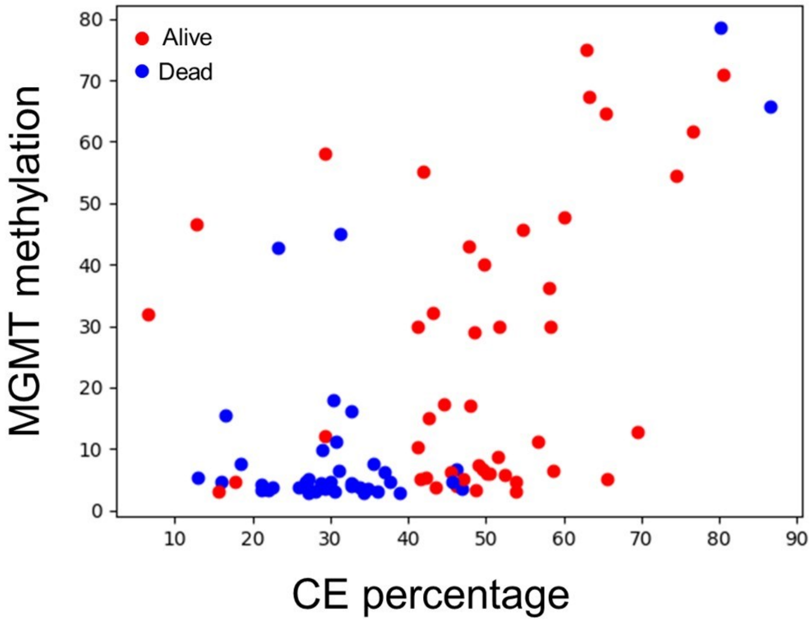


**Supplementary Figure 5. Distribution of CE percentage and MGMT methylation in IDH-wildtype GBM patients stratified by survival status.** Alive (red) and dead (blue) patients are plotted, with CE percentage (x-axis, 0-90) and MGMT methylation (y-axis, 0-80).

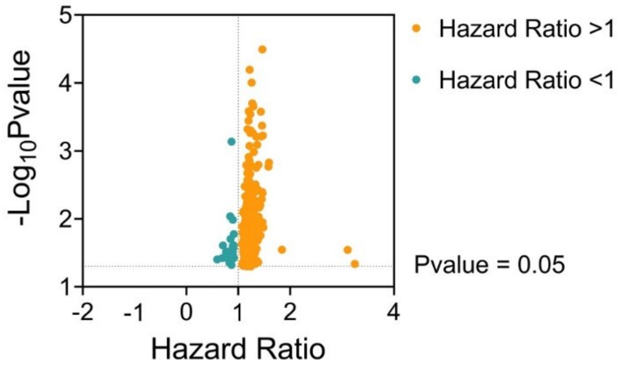


**Supplementary Figure 6. Volcano plot of differentially expressed genes (DEGs) stratified by hazard ratio (HR) in IDH-wildtype GBM prognosis.** DEGs were identified with $P < 0.05$ (horizontal dashed line at $-\log_{10}[P] = 1.3$). Orange dots represent genes with HR > 1, cyan dots denote genes with HR < 1. Axes: x = Hazard Ratio, y = $-\log_{10}$ (Pvalue).

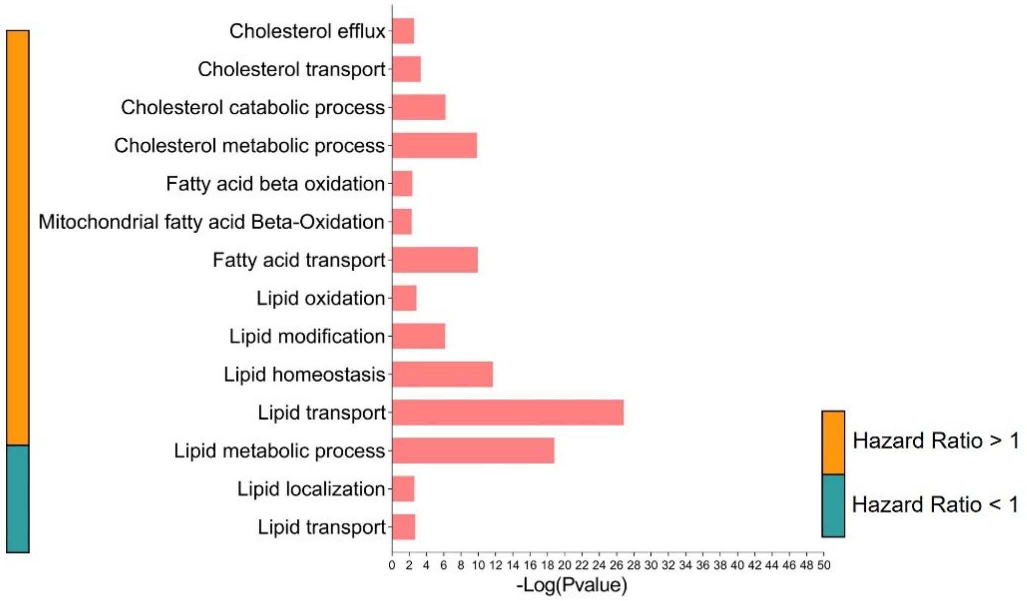


**Supplementary Figure 7. Gene Ontology (GO) pathway enrichment analysis of DEGs stratified by HR in IDH-wildtype GBM prognosis.** Bar length reflects -log$_{10}$ (Pvalue) for pathway enrichment significance; orange bars denote HR > 1 (risk-enhancing), blue bars HR < 1 (protective). Cholesterol efflux, transport, and homeostasis (HR > 1) exhibited the highest significance, correlating with adverse outcomes. Axes: x = -log$_{10}$ (Pvalue), y = GO biological processes.

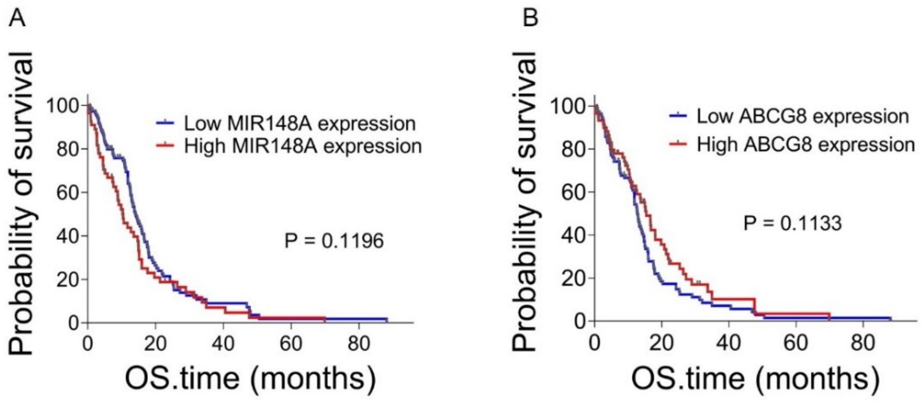


**Supplementary Figure 8. Kaplan-Meier survival analysis of cholesterol metabolism-related genes in IDH-wildtype GBM.** (A) MIR148A and (B) ABCG8 expression levels stratified into low (blue) and high (red) expression groups among 160 patients from the TCGA cohort. Log-rank test Pvalue: MIR148A ($P$ = 0.1196), ABCG8 ($P$ = 0.1133). Survival curves demonstrate no statistically significant difference between groups, suggesting limited prognostic utility for these genes in isolation. Axes: $x$ = overall survival time (months), $y$ = survival probability (%).

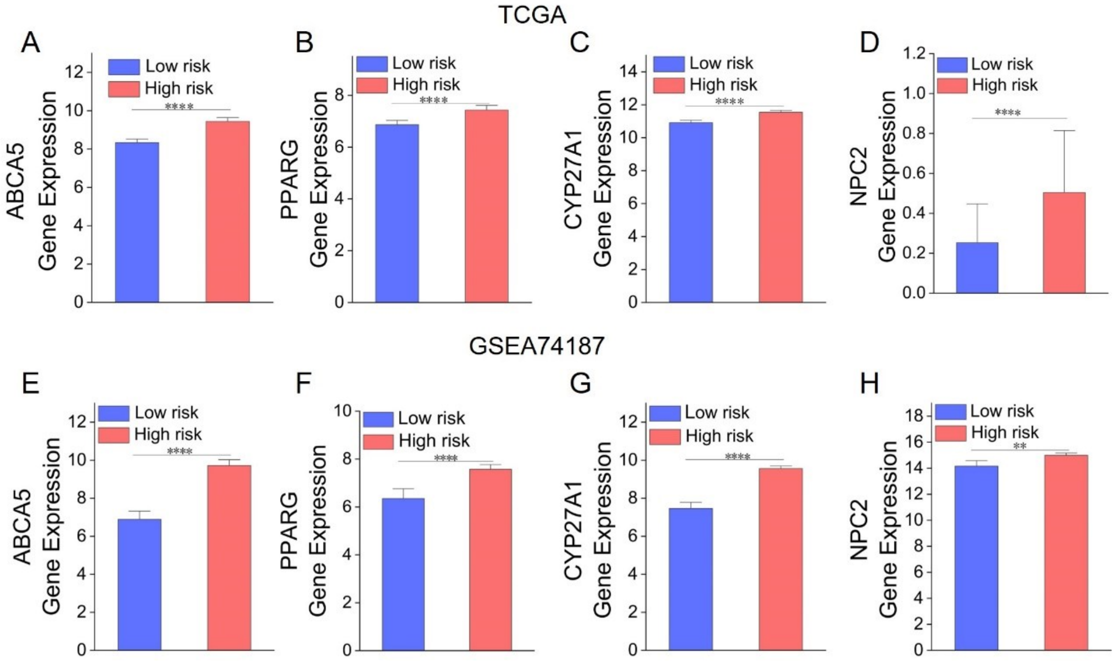


**Supplementary Figure 9. Association between cholesterol metabolism-related gene expression and risk score model stratification in IDH-wildtype GBM.** (A-D) Mean expression levels of (A) ABCA5, (B) PPARG, (C) CYP27A1, and (D) NPC2 in high-risk (red) versus low-risk (blue) groups stratified by the risk score model. Error bars indicate SEM ($n > 10$ per group). One-way ANOVA test: ABCA5 ($P < 0.0005$), CYP27A1 ($P < 0.0005$), PPARG ($P < 0.0005$) and NPC2 ($P < 0.0005$) exhibited significantly lower expression in high-risk groups.

**Supplementary Table 1. Biomarker and clinical information from 107 IDH-wildtype GBM patients.**

| Training cohort | Patient ID | OS (months) | OS status | CE Percentage (%) | MGMT methylation (%) | EGFR amplification | CDKN2A deletion | CDKN2B deletion |
|---|---|---|---|---|---|---|---|---|
| | 1 | 26.13 | 0 | 54.67 | 45.75 | NO | NO | NO |
| | 2 | 24.76 | 0 | 44.54 | 17.25 | YES | YES | NO |
| | 3 | 25.18 | 0 | 53.80 | 4.75 | NO | YES | YES |
| | 4 | 24.10 | 0 | 50.08 | 6.00 | YES | YES | YES |
| | 5 | 24.07 | 0 | 41.18 | 10.25 | YES | YES | YES |
| | 6 | 24.76 | 0 | 29.27 | 12.00 | NO | YES | YES |
| | 7 | 26.14 | 0 | 60.11 | 47.75 | YES | YES | YES |
| | 8 | 24.89 | 0 | 58.30 | 30.00 | YES | YES | YES |
| | 9 | 25.38 | 0 | 12.83 | 46.50 | NO | NO | NO |
| | 10 | 24.53 | 0 | 43.11 | 32.25 | NO | YES | YES |
| | 11 | 26.27 | 0 | 48.64 | 3.25 | YES | YES | YES |
| | 12 | 24.30 | 0 | 80.61 | 71.00 | YES | NO | NO |
| | 13 | 25.87 | 0 | 76.63 | 61.75 | YES | YES | YES |
| | 14 | 24.72 | 0 | 74.50 | 54.50 | NO | YES | YES |
| | 15 | 24.23 | 0 | 6.45 | 32.00 | NO | NO | NO |
| | 16 | 25.02 | 0 | 50.53 | 6.00 | NO | NO | NO |
| | 17 | 26.86 | 0 | 49.52 | 6.75 | NO | NO | NO |
| | 18 | 24.30 | 0 | 41.28 | 30.00 | YES | NO | NO |
| | 19 | 29.85 | 0 | 42.25 | 5.25 | YES | YES | YES |
| | 20 | 29.13 | 0 | 51.57 | 8.75 | NO | NO | NO |
| | 21 | 31.36 | 0 | 46.24 | 4.00 | NO | NO | NO |
| | 22 | 32.32 | 0 | 51.77 | 30.00 | YES | YES | YES |
| | 23 | 25.12 | 0 | 17.77 | 4.75 | YES | NO | NO |
| | 24 | 25.74 | 0 | 41.65 | 5.00 | NO | NO | NO |

| | | | | | | | | |
|---|---|---|---|---|---|---|---|---|
| | 25 | 24.13 | 0 | 15.59 | 3.00 | NO | NO | NO |
| | 26 | 27.32 | 0 | 47.09 | 5.00 | YES | YES | YES |
| | 27 | 26.56 | 0 | 49.12 | 7.25 | NO | YES | YES |
| | 28 | 29.16 | 0 | 58.20 | 36.25 | NO | NO | NO |
| | 29 | 26.99 | 0 | 49.80 | 39.50 | NO | NO | NO |
| | 30 | 30.94 | 0 | 41.90 | 55.25 | NO | YES | YES |
| | 31 | 28.47 | 0 | 29.32 | 57.50 | NO | NO | NO |
| | 32 | 24.93 | 0 | 42.68 | 14.50 | NO | NO | NO |
| | 33 | 29.49 | 0 | 45.56 | 6.25 | NO | YES | YES |
| | 34 | 24.13 | 0 | 43.63 | 3.25 | NO | YES | YES |
| | 35 | 24.16 | 0 | 48.06 | 17.25 | YES | YES | NO |
| | 36 | 25.02 | 0 | 65.62 | 5.00 | YES | YES | YES |
| | 37 | 24.30 | 0 | 63.02 | 75.25 | NO | NO | NO |
| | 38 | 25.94 | 0 | 58.63 | 6.35 | NO | NO | NO |
| | 39 | 25.84 | 0 | 48.54 | 29.00 | NO | NO | NO |
| | 40 | 25.84 | 0 | 53.84 | 3.00 | YES | NO | NO |
| | 41 | 25.38 | 0 | 47.78 | 42.75 | NO | NO | NO |
| | 42 | 30.25 | 0 | 63.36 | 67.25 | NO | NO | NO |
| | 43 | 28.31 | 0 | 56.74 | 11.25 | YES | YES | YES |
| | 44 | 28.01 | 0 | 69.46 | 12.75 | NO | YES | YES |
| | 45 | 26.33 | 0 | 52.40 | 5.75 | NO | YES | YES |
| | 46 | 25.68 | 0 | 65.49 | 64.50 | YES | YES | YES |
| | 47 | 9.21 | 1 | 32.65 | 16.25 | NO | NO | NO |
| | 48 | 17.78 | 1 | 37.65 | 4.75 | YES | NO | NO |
| | 49 | 12.13 | 1 | 16.05 | 4.75 | NO | YES | YES |
| | 50 | 16.21 | 1 | 34.89 | 3.50 | NO | YES | YES |
| | 51 | 8.32 | 1 | 16.51 | 15.50 | NO | NO | NO |
| | 52 | 13.12 | 1 | 27.22 | 5.00 | NO | YES | YES |

| | | | | | | | | |
|---|---|---|---|---|---|---|---|---|
| | 53 | 15.09 | 1 | 86.68 | 65.75 | YES | YES | YES |
| | 54 | 12.82 | 1 | 32.69 | 4.00 | NO | YES | YES |
| | 55 | 14.56 | 1 | 29.30 | 3.50 | YES | YES | YES |
| | 56 | 11.90 | 1 | 28.08 | 3.00 | NO | NO | NO |
| | 57 | 20.62 | 1 | 18.42 | 7.50 | YES | YES | YES |
| | 58 | 12.07 | 1 | 80.11 | 78.50 | NO | YES | YES |
| | 59 | 21.11 | 1 | 22.10 | 3.25 | YES | YES | YES |
| | 60 | 23.11 | 1 | 27.27 | 3.25 | YES | NO | NO |
| | 61 | 4.31 | 1 | 30.55 | 3.00 | NO | YES | YES |
| | 62 | 5.36 | 1 | 12.88 | 5.25 | YES | YES | YES |
| | 63 | 6.01 | 1 | 38.92 | 2.75 | NO | YES | YES |
| | 64 | 6.05 | 1 | 26.62 | 4.00 | NO | YES | YES |
| | 65 | 19.30 | 1 | 21.04 | 3.25 | NO | NO | NO |
| | 66 | 11.93 | 1 | 31.20 | 45.00 | NO | NO | NO |
| | 67 | 6.54 | 1 | 30.00 | 4.75 | YES | YES | YES |
| | 68 | 12.56 | 1 | 28.73 | 4.00 | NO | NO | NO |
| | 69 | 13.74 | 1 | 31.04 | 6.50 | NO | NO | NO |
| | 70 | 18.21 | 1 | 28.82 | 4.50 | YES | YES | YES |
| | 71 | 10.82 | 1 | 28.99 | 9.75 | NO | NO | NO |
| | 72 | 13.25 | 1 | 32.91 | 4.25 | NO | YES | YES |
| | 73 | 20.15 | 1 | 22.62 | 3.75 | NO | YES | YES |
| | 74 | 8.71 | 1 | 27.11 | 2.75 | NO | YES | YES |
| | 75 | 16.44 | 1 | 46.16 | 6.75 | YES | NO | NO |
| | 76 | 12.56 | 1 | 36.07 | 3.00 | NO | YES | YES |
| | 77 | 22.36 | 1 | 33.75 | 3.75 | YES | YES | YES |
| | 78 | 17.98 | 1 | 36.88 | 6.25 | YES | NO | NO |
| | 79 | 23.05 | 1 | 30.65 | 11.25 | NO | YES | YES |
| | 80 | 12.46 | 1 | 21.11 | 4.25 | NO | YES | YES |

| | | | | | | | | |
|---|---|---|---|---|---|---|---|---|
| | 81 | 16.47 | 1 | 46.90 | 3.50 | YES | YES | YES |
| | 82 | 9.24 | 1 | 32.71 | 4.50 | NO | NO | NO |
| | 83 | 11.18 | 1 | 45.71 | 4.75 | NO | NO | NO |
| | 84 | 8.05 | 1 | 30.37 | 18.00 | NO | YES | NO |
| | 85 | 13.81 | 1 | 26.90 | 4.75 | NO | YES | YES |
| | 86 | 14.10 | 1 | 35.60 | 7.50 | YES | YES | YES |
| | 87 | 17.65 | 1 | 23.18 | 42.75 | NO | NO | NO |
| | 88 | 12.92 | 1 | 25.97 | 3.75 | NO | NO | NO |
| | 89 | 6.81 | 1 | 34.37 | 2.75 | YES | YES | YES |
| Testing cohort | 90 | 26.93 | 0 | 42.20 | 3.75 | YES | YES | YES |
| | 91 | 26.66 | 0 | 48.36 | 3.5 | NO | YES | YES |
| | 92 | 26.27 | 0 | 48.73 | 39.5 | NO | YES | YES |
| | 93 | 24.72 | 0 | 47.83 | 33.75 | YES | YES | NO |
| | 94 | 24.20 | 0 | 44.68 | 27 | YES | NO | NO |
| | 95 | 24.04 | 0 | 43.06 | 59.75 | NO | NO | NO |
| | 96 | 19.36 | 1 | 33.75 | 3.00 | YES | YES | YES |
| | 97 | 18.97 | 1 | 46.16 | 20.75 | YES | YES | YES |
| | 98 | 14.43 | 1 | 22.62 | 4.75 | YES | YES | YES |
| | 99 | 16.54 | 1 | 28.24 | 4.25 | NO | YES | YES |
| | 100 | 14.20 | 1 | 31.04 | 8.50 | NO | YES | YES |
| | 101 | 20.71 | 1 | 17.77 | 6025 | NO | NO | YES |
| | 102 | 14.07 | 1 | 34.09 | 5.00 | YES | NO | NO |
| | 103 | 17.03 | 1 | 33.63 | 3.25 | YES | NO | NO |
| | 104 | 20.94 | 1 | 49.80 | 4.00 | YES | NO | NO |
| | 105 | 15.78 | 1 | 32.77 | 49.75 | NO | NO | NO |
| | 106 | 13.91 | 1 | 29.65 | 3.00 | NO | YES | YES |
| | 107 | 17.59 | 1 | 23.20 | 17.50 | NO | NO | NO |

OS status: Two-year Overall Survival (OS), 1 = death, 0 = alive.
